# *In planta* chromatin immunoprecipitation in *Zymoseptoria tritici* reveals chromatin-based regulation of putative effector gene expression

**DOI:** 10.1101/544627

**Authors:** Jessica L. Soyer, Jonathan Grandaubert, Janine Haueisen, Klaas Schotanus, Eva H. Stukenbrock

**Affiliations:** UMR BIOGER, INRA, AgroParisTech, Paris-Saclay University, 78850 Thiverval-Grignon, France; Max Planck Institute for Evolutionary Biology, August-Thienemann-Str. 2, 24306 Plön, and Christian-Albrechts University of Kiel, Am Botanischen Garten 1-9, 24118 Kiel, Germany; Department of Mycology, Fungal Biology and Pathogenicity Unit, Institut Pasteur, INRA, 75015, Paris, France; Departments of Molecular Genetics and Microbiology (MGM), Pharmacology and Cancer Biology, and Medicine, Duke University Medical Center, Durham, North Carolina, United States of America

**Keywords:** Heterochromatin, effectors, gene regulation, ChIP, *Zymoseptoria tritici*, *Triticum aestivum*

## Abstract

During infection, pathogens secrete effectors, key elements of pathogenesis. In several phytopathogenic fungi, synchronous waves of effector genes are expressed during plant infection to manipulate and silence plant defenses. In *Zymoseptoria tritici*, causing septoria leaf blotch of wheat, at least two waves of effector genes are expressed, during the asymptomatic phase and at the switch to necrotrophy. The underlying factors responsible for the fine-tuned regulation of effector gene expression in this pathogen are unknown. Previously, a detailed map of the chromatin structure *in vitro* of *Z. tritici* was generated by chromatin immunoprecipitation followed by high-throughput sequencing (ChIP-seq) targeting histone modifications typical for euchromatin (di-methylation of the lysine 4 of the histone H3, H3K4me2) or heterochromatin (tri-methylation of the lysine 9 and 27 of the histone H3, H3K9me3 and H3K27me3). Based on the hypothesis that changes in the histone modifications contribute to the transcriptional control of pathogenicity-related genes, we tested whether different sets of genes are associated with different histone modifications *in vitro*. We correlated the *in vitro* histone maps with *in planta* transcriptome data and show that genes located in heterochromatic domains *in vitro* are highly up-regulated at the switch toward necrotrophy. We combined our integrated analyses of genomic, transcriptomic and epigenomic data with ChIP-qPCR *in planta* and thereby provide further evidence for the involvement of histone modifications in the transcriptional dynamic of putative pathogenicity-related genes of *Z. tritici*.

## Introduction

Fungi that colonize plant tissues, pathogens as well as mutualists, express effector genes required for their establishment within the plant tissue (Lo Presti *et al*., 2015; Tyler and Rouxel, 2013). Transcriptomic analyzes have shown that effector genes are poorly expressed during axenic growth and strongly induced during host infection with several waves of concerted expression associated to infection stages (e.g. in *Melampsora larici-populina, Colletotrichum* species, *Leptosphaeria maculans;* Gervais *et al*., 2017; Hacquard *et al*., 2012; Lorrain *et al*., 2018; O’Connell *et al*., 2012). The fine-tuned regulation of effector genes is important for successful plant infection by fungi. However, very little is known about the underlying mechanisms that regulate expression of effector genes during different stages of host infection.

Transposable element (TE) rich regions of the genome are often enriched in genes encoding proteinaceous effectors or secondary metabolites (Soyer *et al*., 2015a). In the oilseed rape pathogen *L. maculans*, effector genes expressed during infection of young leaves are enriched in TE-rich regions while effector genes expressed during stem infection are enriched in gene-rich regions of the genome (Gervais *et al*., 2017).

In *L. maculans* and in the distantly related fungi *Epichloe festucae*, a symbiont of the grass *Lolium perenne*, it has been shown that histone modifications play a crucial role in the regulation of genes located in TE-rich regions involved in the establishment of fungus-plant interactions (Chujo and Scott 2014; Soyer *et al*., 2014). These studies support the hypothesis that chromatin-mediated gene regulation represents an efficient strategy to control genes located in the vicinity of TEs.

*Zymoseptoria tritici* causes the septoria tritici blotch disease on wheat (*Triticum aestivum*) (O’Driscoll *et al*., 2014). Infection conferred by *Z. tritici* is initiated by a asymptomatic phase during which the pathogen grows intercellularly. Ten to 12 days post infection, the fungus switches to a necrotrophic stage inducing host cell death. The release of nutrients from dead plant cells allows the formation of pycnidia and the production of asexual spores responsible for new infections. Thus, successful infection of *Z. tritici* involves a lifestyle transition conferred by distinct transcriptional programs (Haueisen *et al*., 2018; Rudd *et al*., 2015; Palma-Guerrero *et al*., 2016). So far, little is known about the regulation of the different infection programs set up by this fungus.

The genome of *Z. tritici* comprises 13 core chromosomes and a variable number of accessory chromosomes showing presence-absence polymorphisms in different isolates (Wittenberg *et al*., 2009; Goodwin *et al*., 2011; Möller *et al*., 2018). TEs represent 18% of the genome with a considerably higher abundance on the accessory chromosomes than on core chromosomes (Dhillon *et al*., 2014; Grandaubert *et al*., 2015). Transcriptomic analyses revealed a dramatic difference in the expression of genes located on core and accessory chromosomes, with the latter showing no or very low expression of most genes *in vitro* as well as *in planta* (Kellner *et al*., 2014; Haueisen *et al*., 2018).

In a previous study, we performed chromatin immunoprecipitation followed by high-throughput sequencing (ChIP-seq) *in vitro* to determine the genome-wide location of histone modifications typical for euchromatin (di-methylation of the lysine 4 of the histone H3, H3K4me2) or with heterochromatin (tri-methylation of the lysines 9 and 27 of H3, H3K9me3 and H3K27me3) (Schotanus *et al*., 2015). In *Z. tritici*, TE-rich regions are enriched with both heterochromatic marks, H3K9me3 and H3K27me3. Furthermore, we confirmed that transcriptional silencing of genes on the accessory chromosomes is due to the strong enrichment with heterochromatic DNA as these chromosomes are almost entirely associated with H3K27me3 (Schotanus *et al*., 2015).

Here, we investigated whether a chromatin-based control plays a role in the transcriptional regulation of pathogenicity-related genes in *Z. tritici*. We combined and integrated genomic, transcriptomic and epigenomic data and performed ChIP followed by quantitative PCR (ChIP-qPCR) analysis of targeted genes during plant infection. We provide evidence for an involvement of H3K9/K27me3 dynamics in the transcriptional regulation of putative pathogenicity genes of *Z. tritici*.

## Results

### Distribution of histone modifications in the coding fraction of the *Zymoseptoria tritici* genome

The ChIP-seq datasets generated from axenic cultures of the isolate Zt09 were analyzed to identify the genes associated with H3K4me2, H3K9me3 or H3K27me3. As previously described, H3K4me2 is mainly associated with gene-rich regions while the heterochromatin mark H3K9me3 is associated with the repetitive regions of the *Z. tritici* genome; H3K27me3 is likewise associated with repetitive sequences, telomeric regions, accessory chromosomes, and with some gene coding sequences on core chromosomes.

We used the genome wide maps of H3K4me2, H3K9me3 and H3K27me3 from *in vitro* growth to distinguish genes either entirely or partially (> 1 bp) associated with each of the three histone modifications. Forty-two percent of the predicted genes in the genome are either associated with H3K4me2 (4,992 out of the 11,754 genes predicted in the genome), 0.75% (89 genes) with H3K9me3, 18.5% (2,179 genes) with H3K27me3 and 1.95% (230 genes) with both H3K9me3 and H3K27me3 (Figure 1A and 1B).

**Figure 1.**
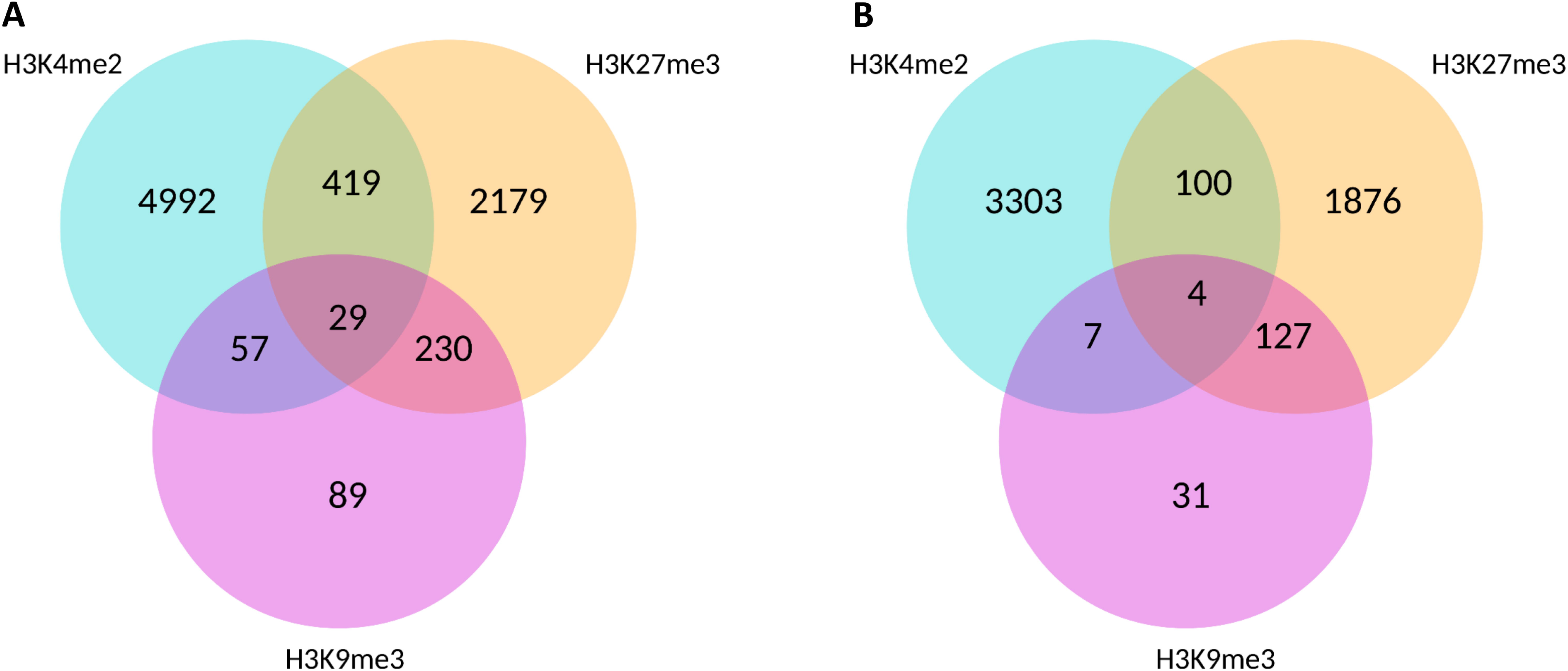
Euchromatic domains encompass more genes than heterochromatin domains *in vitro*. Venn diagrams showing genes associated with the different post-translational histone modifications assessed using ChIP-sequencing in axenic culture. A) Genes at least partially associated with any of the histone modification. B) Genes completely associated with any of the histone modifications.

### Genes located in H3K9me3 and H3K27me3 domains are less conserved and encode proteins putatively involved in virulence

We assessed the correlation of chromatin marks with the distribution of specific-specific genes previously identified in *Z. tritici* (Grandaubert *et al*., 2015). Using comparative genomics of four *Zymoseptoria* species, i.e., *Z. tritici, Zymoseptoria pseudotritici, Zymoseptoria ardabiliae and Zymoseptoria brevis*, we previously identified 1,717 genes unique to *Z. tritici*. We found that these genes are significantly enriched in regions of the genome that are associated with H3K27me3 domains and overlapping H3K9me3/H3K27me3 domains: in total 951 of the *Z. tritici* species-specific genes locate in heterochromatic domains (X^2^ test, *P* < 0.01; Table 1; Table S1). The *Z. tritici* genome encodes 874 putative effector genes and, interestingly, the species-specific genes are enriched in genes encoding putative effectors (177 representing 10.3% of the orphan genes; X^2^ test; *P* < 2.2.10^−6^) of which 82 locate in H3K9me3 and / or H3K27me3 domains. In the *Z. tritici* reference genome, 6,329 predicted genes could not be assigned to a protein function (i.e. no homology with proteins predicted in other organisms and/or no protein domain included in databases). Hereinafter, we refer to these as genes of “unknown function”. We found a significant enrichment of genes of unknown function in the heterochromatic domains enriched with H3K27me3 or enriched with both H3K9me3 and H3K27me3 (X^2^ test, *P* < 0.01; Table 1; Table S2). Contrarily, genes associated with H3K4me2 were deprived of genes of unknown function compared to the rest of the genome (X^2^ test, *P* < 0.01; Table S2). Taken together, these data suggest that heterochromatic DNA associates with less conserved genes, including many predicted to be secreted and involved in host-pathogen interactions.

**Table 1.** Number of genes in a given category according to their location in a H3K4me2, H3K9me3, H3K27me3 or H3K9me3/H3K27me3 domain *in vitro*.

|  | genome |  | H3K4me2-domains<sup>f</sup> |  | H3K9me3-domains<sup>f</sup> |  | H3K27me3-domains<sup>f</sup> |  | H3K9me3+H3K27me3-domains<sup>f</sup> |  |
| --- | --- | --- | --- | --- | --- | --- | --- | --- | --- | --- |
|  | nb. in the entire genome | nb. in the core genome | nb. in the entire genome | nb. in the core genome | nb. in the entire genome | nb. in the core genome | nb. in the entire genome | nb. in the core genome | nb. in the entire genome | nb. in the core genome |
| total number of genes | 11754 | 11111 | 4992 | 4991 | 89 | 89 | 2179 | 1661 | 230 | 171 |
| unknown<sup>a</sup> | 6329 | 5699 | 1944 | 1943 | 50 | 50 | 1734 | 1223 | 193 | 152 |
| PKS/NRPS | 20 | 20 | 1 | 1 | 0 | 0 | 10 | 10 | 0 | 0 |
| CAZymes | 227 | 227 | 58 | 58 | 4 | 4 | 32 | 32 | 3 | 3 |
| genes < 2 Kb Tes<sup>b</sup> | 1723 | 1464 | 369 | 368 | 83 | 83 | 545 | 358 | 202 | 154 |
| orphan genes | 1717 | 1221 | 177 | 177 | 21 | 21 | 794 | 392 | 136 | 91 |
| genes expressed at 4 dpi vs axenic<sup>c</sup> | 559 | 546 | 92 | 92 | 4 | 4 | 149 | 138 | 9 | 9 |
| genes expressed at 13 dpi vs axenic<sup>c</sup> | 886 | 778 | 98 | 98 | 10 | 10 | 355 | 253 | 20 | 17 |
| genes expressed at 13 dpi vs 4 dpi<sup>c</sup> | 729 | 577 | 53 | 53 | 10 | 10 | 368 | 224 | 27 | 20 |
| genes up-regulated at 4 dpi vs. *in vitro*<sup>d</sup> | 784 | 775 | 290 | 290 | 6 | 6 | 118 | 112 | 10 | 10 |
| genes down-regulated at 4 dpi vs. *in vitro*<sup>e</sup> | 1027 | 1001 | 379 | 379 | 3 | 3 | 179 | 156 | 23 | 21 |
| genes up-regulated at 13 dpi vs. 4 dpi<sup>d</sup> | 1773 | 1630 | 388 | 388 | 12 | 12 | 507 | 390 | 38 | 32 |
| genes down-regulated at 13 dpi vs. 4 dpi<sup>e</sup> | 798 | 796 | 385 | 385 | 5 | 5 | 81 | 80 | 12 | 11 |

<sup>a</sup>“unknown” genes are genes encoding predicted or hypothetical proteins;
<sup>b</sup>Genes located up to 2 kb upstream or downstream to a transposable element sequence;
<sup>c</sup>Genes specifically expressed at 4 or 13 days post infection (dpi) compared to axenic culture or 4 dpi when RPKM $\geq 2$ at the time point and RPKM $\leq 2$ during the axenic culture or at 4 dpi;
<sup>d</sup>Up-regulated genes are genes with Log2 Fold Change (RPKM) $\geq 1$ and FDR $\leq 0.001$ in a given condition compared to the other;
<sup>e</sup>Down-regulated genes are genes with Log2 Fold Change (RPKM) $\leq -1$ and FDR $\leq 0.001$ in a given condition compared to the other;
<sup>f</sup>Genes located in a H3K4me2, H3K9me3, H3K27me3 or H3K9me3/H3K27me3 domain *in vitro*.

### H3K9me3 and H3K27me3 are associated with genes encoding putative pathogenicity determinants

Specific types of proteins are often associated with pathogenicity in fungi (for example proteinaceous effectors and proteins involved in secondary metabolite syntheses; Tyler and Rouxel, 2013). In some species, such as Aspergilli, secondary metabolite-encoding genes were found to be enriched in subtelomeric regions of the genome or near TEs (Gacek and Strauss, 2012). Furthermore, a chromatin-based regulation of these genes by post-translational histone modifications was demonstrated for gene clusters encoding polyketide synthase-like (PKS) proteins and non-ribosomal peptide synthase (NRPS) gene clusters, e.g. in the endophyte *E. festucae* (Chujo and Scott, 2014).

We assessed whether PKS and NRPS encoding genes are associated with the two heterochromatin marks studied here. In total, 2,498 predicted genes (21% of all genes) locate in heterochromatic domains during *in vitro* growth of *Z. tritici*. Among these, we identified several genes involved in secondary metabolism and detoxification, i.e. with a predicted function such as cytochrome P450, polyketide synthase or Major Facilitator Superfamily (MFS) transporters. More precisely, 11 genes of the *Z. tritici* genome were annotated as PKS and nine as NRPS encoding genes (Ohm *et al*., 2012; Grandaubert *et al*., 2015). PKS/NRPS encoding genes are significantly enriched in H3K27me3 domains as identified during *in vitro* growth of the fungus (X^2^ test, *P* < 0.01, Table 1; Table S1).

We next used a PFAM enrichment analysis (Finn *et al*., 2014) to assess if genes encoding proteins involved in biosynthetic activities were also enriched in heterochromatic domains of the *Z. tritici* genome. We found that H3K27me3 and H3K9me3 domains are significantly enriched in genes involved in biosynthetic activities including genes encoding Rad51 PFAM domains known to be involved in DNA repair processes and genes encoding reverse transcriptases and transposases. Furthermore, genes located in H3K27me3 domains are enriched in PFAM protein families involved in secondary metabolism (Tables S3-6; FDR < 0.01).

We also analyzed the location of 227 genes predicted to encode Carbohydrate-Active Enzymes (CAZymes) in the *Z. tritici* genome. Contrary to secondary metabolite genes, there is no significant enrichment of the CAZyme genes in heterochromatic domains (X^2^ test, *P* > 0.01; Table 1; Table S2).

In the pathogenic fungus *L. maculans*, effector genes showing stage-specific expression profiles during infection of oilseed rape (*Brassica napus*), are enriched in GC-poor or TE-rich genomic regions (Gervais *et al*., 2017). Interestingly, the silencing of genes located in TE-rich regions, *in vitro*, involves a chromatin-based control via H3K9me3 (Soyer *et al*., 2014). We addressed whether TE-rich genomic regions and heterochromatic domains of the *Z. tritici* genome likewise are enriched in putative effector genes. The 874 predicted effector genes represent eight percent of all predicted genes in *Z. tritici* and only nine on the accessory chromosomes (1.04% of all genes predicted on the accessory chromosomes). We here refined our analyses to address the association of the 865 putative effector genes of the core chromosomes and TEs. We considered a gene to be associated with TE regions if the gene locates within 2 kb distance of a TE. Assigning this criterion, we confirm that TE-rich regions of core chromosomes are enriched in putative effector genes (185 putative effector genes corresponding to 12.6% of the TE associated genes; X^2^ test, *P* < 0.01; Table 2; Table S7; Figure 2). We also found a significant enrichment of putative effector genes in heterochromatic domains associated with TEs or not (H3K9me3, H3K27me3 and H3K9me3/H3K27me3 overlapping domains; Table 2; Table S7) while euchromatic domains are deprived of putative effector genes (X^2^ test, *P* < 0.01; Table S7). Altogether, our analysis suggests that genes encoding putative pathogenicity-determinants are non-randomly distributed in the genome of *Z. tritici* and in particular enriched in H3K27me3 and H3K9me3 domains *in vitro*.

**Figure 2.**
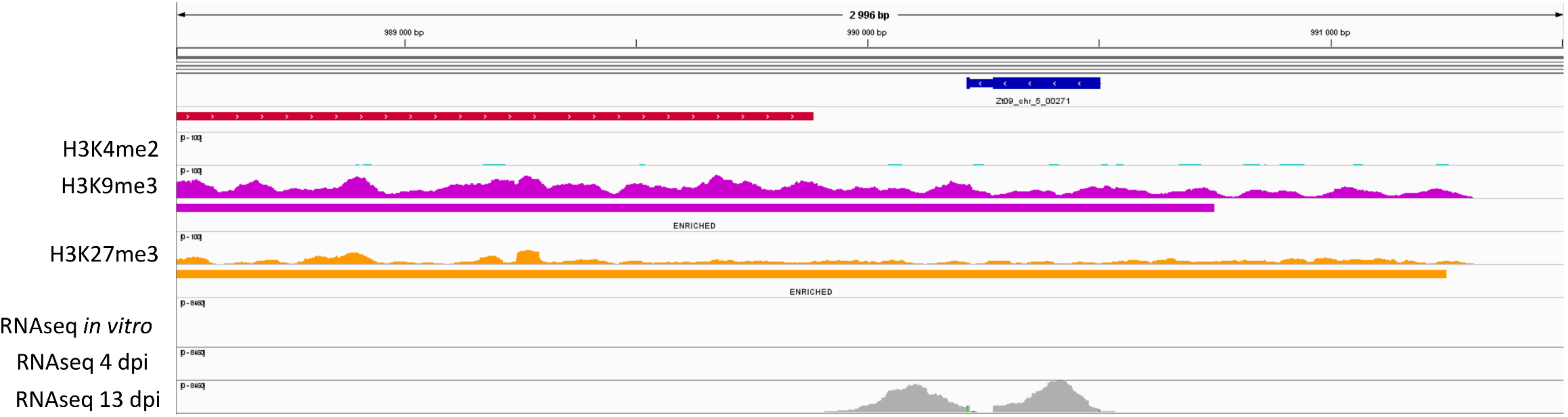
Heterochromatic domains are enriched with putative effector genes that are up-regulated during the switch towards necrotrophic growth on wheat. Example from a section of chromosome 5, harboring the putative effector gene ID Zt09_chr_5_00271. Regions enriched in different histone modifications were identified using ChIP-seq with antibodies against H3K4me2 (light blue), H3K9me3 (purple), H3K27me3 (orange) and compared to the location of genomic features (coding sequences, dark blue; transposable elements, red) and to reads obtained by RNA-seq in axenic culture, 4 days post-infection (dpi) and 13 dpi.

**Table 2.** Number of putative effector genes located in vicinity to a transposable element or in a H3K4me2, H3K9me3, H3K27me3 and H3K9me3/H3K27me3 domain *in vitro*.

|  | core genome | TE-associated genes<sup>c</sup> | H3K4me2-domains<sup>d</sup> | H3K9me3-domains<sup>d</sup> | H3K27me3-domains<sup>d</sup> | H3K9+H3K27me3-domains<sup>d</sup> |
| --- | --- | --- | --- | --- | --- | --- |
| total number of genes | 11111 | 1464 | 4991 | 89 | 1661 | 171 |
| putative effector genes | 865 | 185 | 168 | 15 | 205 | 26 |
| putative effector genes up-regulated at 4 dpi vs axenic<sup>a</sup> | 114 | 14 | 24 | 3 | 22 | 1 |
| putative effector genes down-regulated at 4 dpi vs axenic<sup>b</sup> | 107 | 32 | 23 | 0 | 29 | 6 |
| putative effector genes up-regulated at 13 dpi vs 4 dpi<sup>a</sup> | 309 | 84 | 42 | 4 | 87 | 12 |
| putative effector genes down-regulated at 13 dpi vs 4 dpi<sup>b</sup> | 74 | 10 | 22 | 1 | 10 | 1 |

Only putative effector genes located in the core genome of *Z. tritici* are shown here.
<sup>a</sup>Up-regulated genes are genes with Log2 Fold Change (RPKM) $\geq 1$ and FDR $\leq 0.001$ in a given condition compared to the other;
<sup>b</sup>Down-regulated genes are genes with Log2 Fold Change (RPKM) $\leq -1$ and FDR $\leq 0.001$ in a given condition compared to the other;
<sup>c</sup>Genes located up to 2 kb upstream or downstream to a transposable element sequence;
<sup>d</sup>Genes located in a H3K4me2, H3K9me3, H3K27me3 or H3K9me3/H3K27me3 *in vitro*.

### *In vitro* heterochromatic domains are enriched with genes up-regulated upon plant infection

As genes predicted as putative pathogenicity determinants (PKS, NRPS, effector genes) appear not to be randomly located in the genome of *Z. tritici*, we hypothesize that regulation of the expression of these genes involves a chromatin-based control as shown for effectors in *L. maculans* (Soyer *et al*., 2014). We therefore assessed the correlation of the genome-wide expression patterns in the *Z. tritici* isolate Zt09 with the distribution of genome-wide map of H3K4me2, H3K9me3 and H3K27me3 histone modifications observed during *in vitro* growth (Schotanus *et al*., 2015). To this end, we processed transcriptomic data generated from axenic cultures and infected plant material from previous studies (Kellner *et al*., 2014, Haueisen *et al*., 2018).

During axenic growth, 9,412 genes are expressed with RPKM ≥ 2. Ninety-eight percent (3,237) of the genes for which sequence is entirely associated with H3K4me2, 58% and 39% of genes (i.e. 18 and 737 genes) associated with H3K9me3 or H3K27me3 respectively were expressed *in vitro* (RPKM ≥ 2). These patterns confirm that H3K4me2 is almost systematically associated with transcriptional activity while H3K9me3 and H3K27me3 represent repressive histone modifications (Wilcoxon-test, *P* < 0.05; Figure 3).

**Figure 3.**
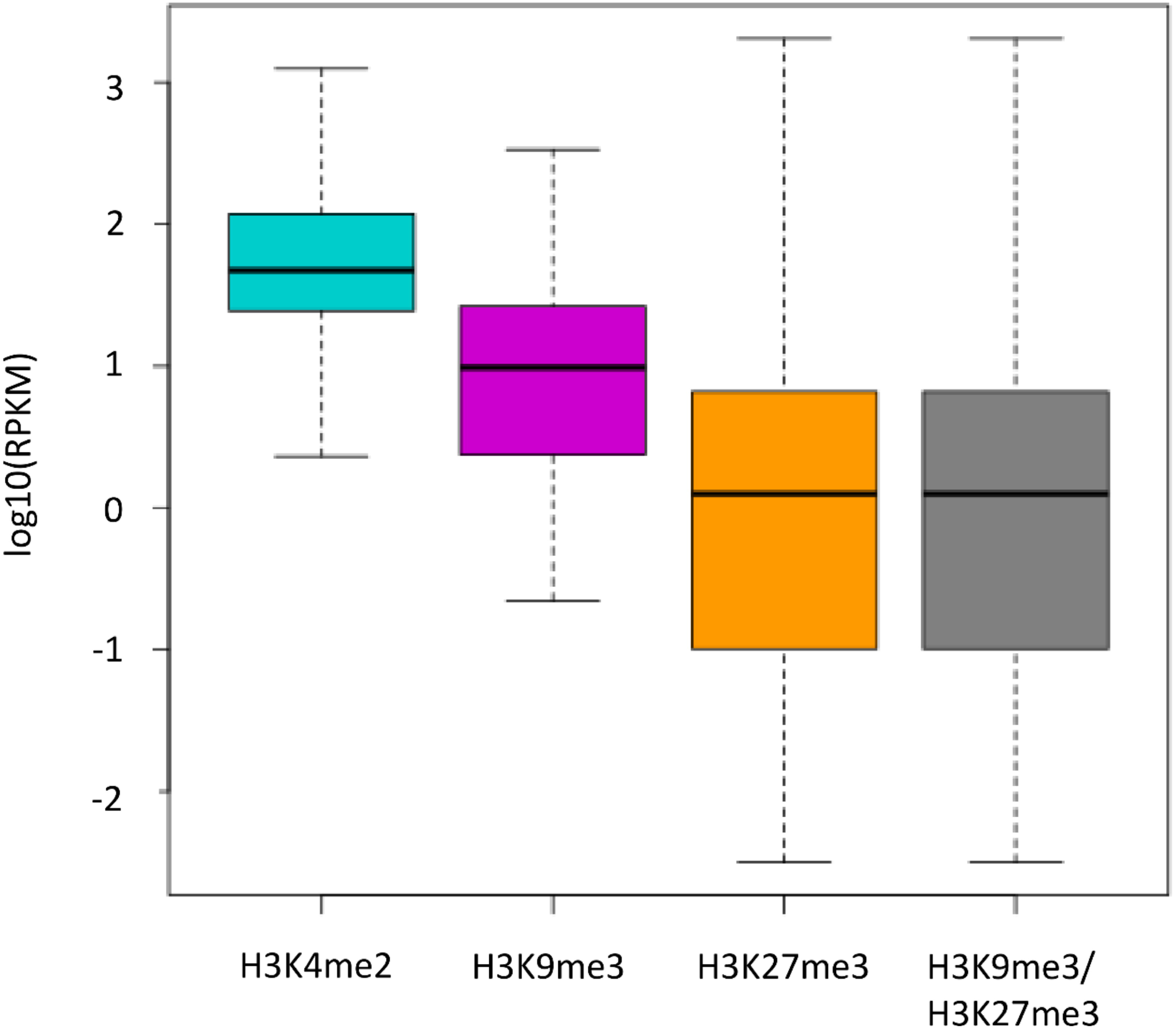
Genes associated with euchromatin are expressed while genes associated with heterochromatin are repressed. Expression assessed during axenic culture using RNA-sequencing. Boxplot of the log10(RPKM) of the genes totally associated with any of the histone modification.

We then compared the transcriptomic profiles of Zt09 genes during host infection and assessed genome wide expression patterns at 4 and 13 days post inoculation (dpi), i.e. during early asymptomatic host infection and at the transition to necrotrophic growth, respectively. We analyzed 1) the number and distribution of genes silenced during *in vitro* growth and induced during plant infection and 2) the number and distribution of genes significantly up- or down-regulated *in planta* compared to axenic culture (Figure 4).

**Figure 4.**
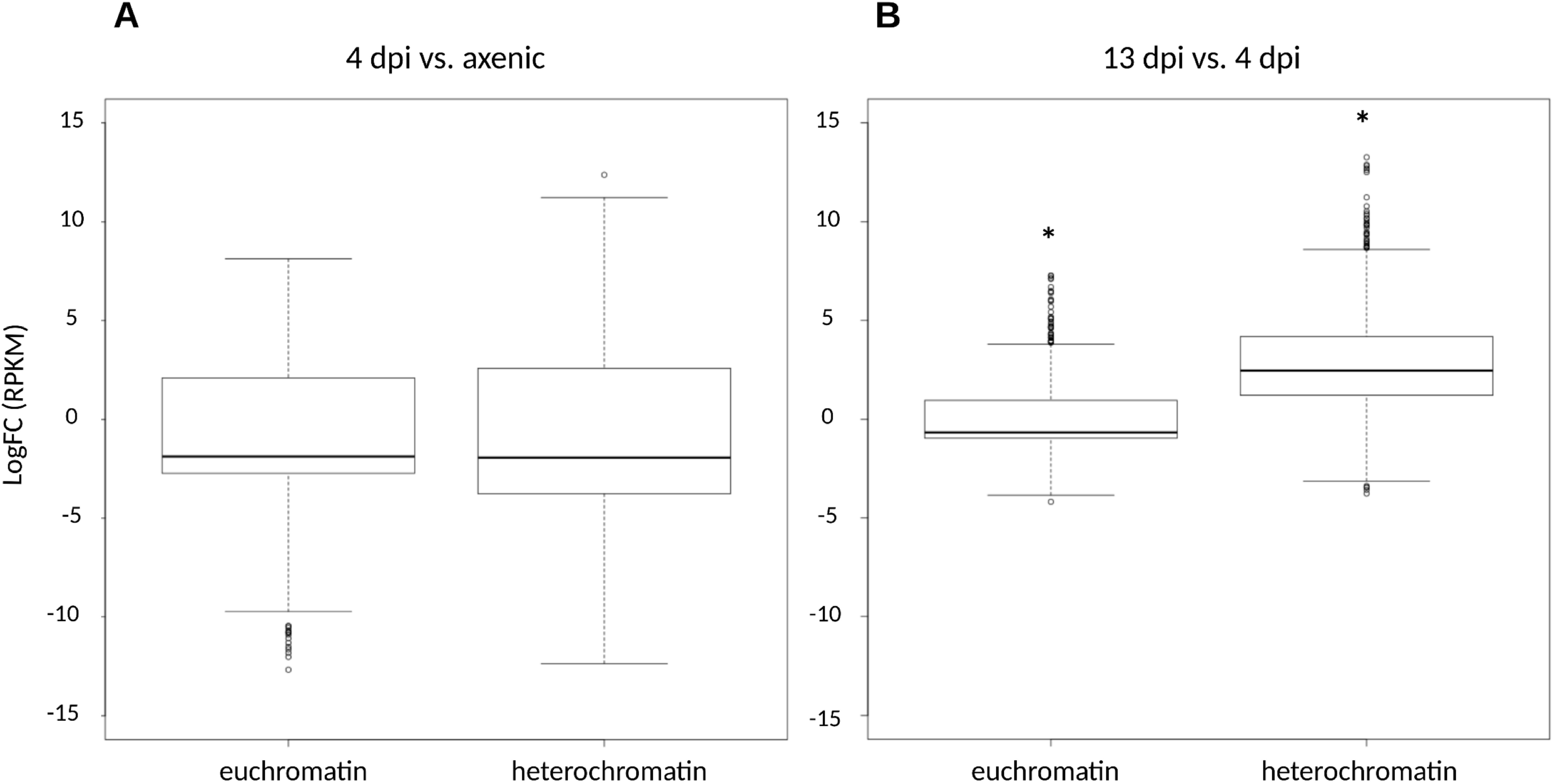
Genes associated with euchromatin and heterochromatin *in vitro* exibit different expression profiles *in planta*. Boxplot of the log_10_ Fold Change (RPKM) of the genes located in a euchromatic domain *in vitro* (i.e. H3K4me2) or a heterochromatic domain *in vitro* (i.e. H3K9me3, H3K27me3 or H3K9me3/H3K27me3). Expression assessed during axenic culture and at 4 and 13 days post infection (dpi) using RNA-sequencing; genes differentially expressed at 4 dpi vs. axenic culture (A) or 13 dpi vs. 4 dpi (B). *: Wilcoxon-test, *P* < 2.2.10^−16^: comparison of heterochromatic-associated genes between 4 and 13 dpi.

We first analyzed genes specifically induced at 4 or 13 dpi compared to axenic culture (i.e. RPKM ≤ 2 for the axenic culture and RPKM ≥ 2 at 4 or 13 dpi). Using these criteria we found 559 genes induced at 4 dpi, including 92 genes that locate in an *in vitro* H3K4me2 domain and 149 in an *in vitro* H3K27me3 domain (the H3K4me2 domains comprise a total of 4,992 genes and H3K27me3 a total of 2,179 genes) (Table 1; Table S2). This pattern shows that H3K27me3 domains are enriched in genes specifically expressed at 4 dpi compared to axenic culture (X^2^ test, *P* < 0.01; Table 1; Table S2). At 13 dpi, 886 genes, not expressed *in vitro*, are induced (i.e. RPKM ≤ 2 for the axenic culture and RPKM ≥ 2 at 13 dpi), including 98 genes located in an *in vitro* H3K4me2 domain and 355 genes located in an *in vitro* H3K27me3 domain. For this later stage of infection, as for 4 dpi, we also found that genes specifically expressed at 13 dpi compared to axenic culture are enriched in genes associated with *in vitro* H3K27me3 domains (X^2^ test, *P* < 0.01; Table 1; Table S2).

We next analyzed genes up-regulated *in planta* compared to the axenic culture (log2 Fold Change (RPKM) ≥ 1 and FDR ≤ 0.001). At 4 dpi, 784 genes were significantly up-regulated compared to the axenic growth. These up-regulated genes are significantly enriched in genes encoding effector candidates (114 putative effector genes of all up-regulated genes at 4 dpi; X^2^ test, *P* < 0.01; Table 2; Table S7). However, during early infection, up-regulated genes are not significantly associated with any of the histone modifications investigated here. At 4 dpi, there is no effect of the *in vitro* location in H3K4me2, H3K9me3 or H3K27me3 on the up- or down-regulation of genes (Table 1; Table S2; Figure 4A). At 13 dpi, 1,773 genes were specifically up-regulated compared to 4 dpi, including 388 genes and 507 genes located respectively in a H3K4me2 and in a H3K27me3 domain *in vitro*. As for 4 dpi, up-regulated genes were significantly enriched in effector genes (309 putative effector genes; X^2^ test, *P* < 0.01; Table 2, Table S7). At 13 dpi, transcription of the genes located in an *in vitro* heterochromatic domain is significantly different from genes located in an *in vitro* euchromatic domain (Wilcoxon test, *P* < 2.2.10^−16^; Figure 4B). We observe that genes showing low, or no, expression *in vitro* and associated with a histone modification typical for heterochromatin in this growing condition are significantly higher expressed at the switch to necrotrophy than genes associated *in vitro* with euchromatin (Table 1; Table S2). Up-regulated effector genes were likewise enriched in genes located in the vicinity of TEs (i.e. distance ≤ 2 kb) and H3K27me3 *in vitro* (Table 2; Table S7). Taken together, these data show that although sets of putative effector genes were highly expressed both 4 dpi and 13 dpi, the histone modifications associated with the genes possibly influences their expression only at the switch to necrotrophy.

### *KMT1, KMT6* or *KMT1/KMT6* deletions deregulate expression of pathogenicity-related genes

In *Z. tritici, kmt1, kmt6* and *kmt1/kmt6* mutants were generated and gene expression was analyzed, *in vitro*, using RNA-seq (Moeller *et al*., 2018). We processed these data to address whether KMT1 and / or KMT6 may influence pathogenesis-related genes.

Among the 100 most up-regulated genes in respectively the *kmt1, kmt6* and *kmt1/kmt6* mutants, 59, 36 and 32% are over-expressed at 13 dpi compared to axenic culture in the WT strain suggesting that both histone methyltransferases are involved in regulation of genes highly expressed in the WT strain upon infection. We analyzed whether certain categories of genes were significantly enriched among the deregulated genes in the mutant background (Table 3). *KMT1* deletion induces the reorganization of H3K27me3 modification with a relocation of this modification to H3K9me3 regions (Moeller *et al*., 2018). Hence, genes located in H3K9me3 are not significantly induced because of H3K9me3 loss as they may still be repressed by the H3K27me3 modification (Table 3). However, consistently with the relocation of H3K27me3, genes located in a H3K27me3 domain in the WT are significantly induced in the *kmt1* mutant (Table 3). We identified that deregulated genes in the *kmt1, kmt6* or *kmt1/kmt6* mutants were significantly enriched in SP-genes (i.e. 220 putative effectors within the 2,110 deregulated genes) and genes associated with TE sequences while no significant association of the CAZymes encoding genes was observed (Table 3; X^2^ test, *P* < 0.01). Particularly, the 972 genes up-regulated in at least one of the mutants include 118 putative effector genes. Deletions of *KMT1* and *KMT6* also significantly influenced expression of genes otherwise found to be induced during infection in the WT strain (Table 3). Genes up-regulated at 4 dpi compared to axenic culture, or 13 dpi compared to 4 dpi, in the WT strain are also significantly enriched in the deregulated genes in the different mutants (231 out of the 784 genes up-regulated at 4 dpi and 646 out of the 1,773 genes up-regulated at 13 dpi are deregulated in the mutants).

**Table 3.** Influence of *kmt1, kmt6* or *kmt1/kmt6* deletions on gene expression.

Number of genes deregulated due to *KMT1*, *KMT6* or *KMT1/KMT6* deletions, *in vitro*.
|  | genome |  | *Δkmt1<sup>h</sup>* |  |  |  | *Δkmt6<sup>h</sup>* |  |  |  |
| --- | --- | --- | --- | --- | --- | --- | --- | --- | --- | --- |
|  | nb. in the genome | proportion | up-regulated |  | down-regulated |  | up-regulated |  | down-regulated |  |
|  |  |  | nb. in the genome | proportion | nb. in the genome | proportion | nb. in the genome | proportion | nb. in the genome | proportion |
| total number of genes | 11754 |  | 579 |  | 284 |  | 274 |  | 481 |  |
| H3K4me2<sup>a</sup> | 4992 | 0.425 | 116 | **0.200** | 71 | **0.250** | 92 | **0.336** | 194 | 0.403 |
| H3K9me3<sup>a</sup> | 89 | 0.008 | 4 | 0.007 | 8 | 0.028 | 0 |  | 2 | 0.004 |
| H3K27me3<sup>a</sup> | 2179 | 0.185 | 168 | 0.290 | 57 | 0.201 | 116 | 0.423 | 63 | **0.131** |
| H3K9/K27me3<sup>a</sup> | 230 | 0.020 | 20 | 0.035 | 16 | 0.056 | 23 | 0.084 | 5 | 0.010 |
| unknown<sup>b</sup> | 6329 | 0.538 | 399 | 0.689 | 117 | 0.412 | 175 | 0.639 | 175 | **0.364** |
| CAZymes | 227 | 0.019 | 10 | 0.017 | 9 | 0.032 | 7 | 0.026 | 7 | 0.015 |
| putative effectors | 874 | 0.074 | 88 | 0.152 | 56 | 0.197 | 24 | 0.088 | 59 | 0.123 |
| genes < 2 Kb Tes<sup>c</sup> | 1723 | 0.147 | 112 | 0.193 | 83 | 0.292 | 79 | 0.288 | 64 | 0.133 |
| orphan genes | 1717 | 0.146 | 101 | 0.174 | 33 | 0.116 | 88 | 0.321 | 27 | **0.056** |
| genes expressed at 4 dpi vs axenic<sup>d</sup> | 559 | 0.048 | 55 | 0.095 | 0 | NA | 3 | 0.011 | 7 | **0.015** |
| genes expressed at 13 dpi vs axenic<sup>d</sup> | 886 | 0.075 | 121 | 0.209 | 2 | 0.007 | 25 | 0.091 | 9 | **0.019** |
| genes expressed at 13 dpi vs 4 dpi<sup>d</sup> | 729 | 0.062 | 105 | 0.181 | 12 | 0.042 | 44 | 0.161 | 21 | 0.044 |
| genes up-regulated at 4 dpi vs. *in vitro*<sup>e</sup> | 784 | 0.067 | 104 | 0.180 | 22 | 0.077 | 11 | 0.040 | 72 | 0.150 |
| genes down-regulated at 4 dpi vs. *in vitro*<sup>f</sup> | 1027 | 0.087 | 61 | 0.105 | 55 | 0.194 | 34 | 0.124 | 120 | 0.249 |
| genes up-regulated at 13 dpi vs. 4 dpi<sup>e</sup> | 1773 | 0.151 | 276 | 0.477 | 78 | 0.275 | 92 | 0.336 | 120 | 0.249 |
| genes down-regulated at 13 dpi vs. 4 dpi<sup>f</sup> | 798 | 0.068 | 48 | 0.083 | 61 | 0.215 | 15 | 0.055 | 113 | 0.235 |

| *Δkmt1/kmt6<sup>h</sup>* |  |  |  |  |
| --- | --- | --- | --- | --- |
|  | up-regulated |  | down-regulated |  |
|  | nb. in the genome | proportion | nb. in the genome | proportion |
| total number of genes | 558 |  | 455 |  |
| H3K4me2<sup>a</sup> | 72 | **0.129** | 147 | **0.323** |
| H3K9me3<sup>a</sup> | 8 | 0.014 | 10 | 0.022 |
| H3K27me3<sup>a</sup> | 248 | 0.444 | 69 | 0.152 |
| H3K9/K27me3<sup>a</sup> | 29 | 0.052 | 22 | 0.048 |
| unknown<sup>b</sup> | 420 | 0.753 | 157 | **0.345** |
| CAZymes | 7 | 0.013 | 10 | 0.022 |
| putative effectors | 65 | 0.116 | 62 | 0.136 |
| genes < 2 Kb Tes<sup>c</sup> | 138 | 0.247 | 125 | 0.275 |
| orphan genes | 166 | 0.297 | 50 | 0.110 |
| genes expressed at 4 dpi vs axenic<sup>d</sup> | 62 | 0.111 | 0 | NA |
| genes expressed at 13 dpi vs axenic<sup>d</sup> | 137 | 0.246 | 2 | **0.004** |
| genes expressed at 13 dpi vs 4 dpi<sup>d</sup> | 119 | 0.213 | 18 | 0.040 |
| genes up-regulated at 4 dpi vs. *in vitro*<sup>e</sup> | 94 | 0.168 | 25 | 0.055 |
| genes down-regulated at 4 dpi vs. *in vitro*<sup>f</sup> | 33 | 0.059 | 123 | 0.270 |
| genes up-regulated at 13 dpi vs. 4 dpi<sup>e</sup> | 229 | 0.410 | 117 | 0.257 |
| genes down-regulated at 13 dpi vs. 4 dpi<sup>f</sup> | 38 | 0.068 | 73 | 0.160 |

<sup>a</sup>Genes located in a H3K4me2, H3K9me3, H3K27me3 or H3K9me3, H3K27me3 *in vitro*;
<sup>b</sup>“unknown” genes are genes encoding predicted or hypothetical proteins;
<sup>c</sup>Genes located up to 2 kb upstream or downstream to a transposable element sequence;
<sup>d</sup>Genes specifically expressed at 4 or 13 days post infection (dpi) compared to axenic culture or 4 dpi when RPKM $\geq 2$ at the time point and RPKM $\leq 2$ during the axenic culture or at 4 dpi;
<sup>e</sup>Up-regulated genes are genes with Log2 Fold Change (RPKM) $\geq 1$ and FDR $\leq 0.001$ in a given condition compared to the other;
<sup>f</sup>Down-regulated genes are genes with Log2 Fold Change (RPKM) $\leq -1$ and FDR $\leq 0.001$ in a given condition compared to the other;
<sup>h</sup>Genes deregulated in *KMT1*, *KMT6* or *KMT1/KMT6* deletions background, Log2 Fold Change (RPKM) $\leq -1$ or $\geq 1$ and FDR $\leq 0.001$ .
Bold: the given domain contains significantly less genes of the given category compared to the rest of the genome; grey: the given domain is enriched for the category of genes compared to the rest of the genome.

We analyzed the expression of a few randomly selected genes based on their location in a heterochromatin domain *in vitro*, by RT-qPCR, in *kmt1, kmt6* and *kmt1/kmt6* deletion mutants. Some of these genes are putative effector genes or orphan genes and some are up-regulated in the WT at 13 dpi compared to early infection (4 dpi) in wheat (Table 4). One gene located in a H3K27me3 domain was not influenced by none of the deletion (gene ID Zt09_chr_3_00231) while all other genes had their expression up-regulated due to the deletion of at least *KMT1, KMT6* or both.

**Table 4.** Expression of selected genes associated with heterochromatin *in vitro*, in the *KMT1, KMT6* and *KMT1/KMT6* deletion mutants.

|  |  |  |  |  |  |  | *in planta* expression<sup>d</sup> | relative expression vs. WT<sup>e</sup> |  |  |
| --- | --- | --- | --- | --- | --- | --- | --- | --- | --- | --- |
| ID | orphan<sup>a</sup> | TE<sup>b</sup> | SP<sup>a</sup> | H3K4me2<sup>c</sup> | H3K9me3<sup>c</sup> | H3K27me3<sup>c</sup> | up 13 vs 4 dpi | $\Delta kmt1$ | $\Delta kmt6$ | $\Delta kmt1/kmt6$ |
| Zt09_chr_2_00407 | - | yes |  | - | yes |  | - | 2 | 3 | 7 |
| Zt09_chr_2_00483 | yes | - | - | - | - | yes | - |  | 3 | 2-8 |
| Zt09_chr_3_00231 | yes | yes | yes | - | - | yes | 6 | - | - | - |
| Zt09_chr_4_00279 | - | yes | yes | - | yes |  | 6 | - | 3-5 | - |
| Zt09_chr_8_00723 | yes | yes |  | - |  |  | 3 | 2-5 | 200 | 20 |
| Zt09_chr_9_00004 | - | yes |  | - | yes | yes | 3 | 6-16 | 65-90 | ~9 |
| Zt09_chr_9_00005 | - | yes | yes | - | yes | yes | 8 | - | 40 | ~3 |
| Zt09_chr_11_00525 | - | yes |  | - | yes | yes | 6 | 20-69 | 200-1000 | 140 |
| Zt09_chr_13_00020 | - | - | - | - | - | yes | 10 | 1-4 | 28-36 | 6-9 |

<sup>a</sup>Genes located up to 2 kb upstream or downstream to a transposable element sequence;
<sup>b</sup>Genes located in a H3K4me2, H3K9me3, H3K27me3 or H3K9me3, H3K27me3 *in vitro*;
<sup>c</sup>Genes significantly up-regulated 13 days post-infection (dpi) compared to 4 dpi;
<sup>d</sup>Gene expression relative to the WT strain and to the GAPDH encoding gene.

### Up-regulation of effector candidate genes *in planta* is associated with changes in chromatin structure

The analysis of *in vitro* ChIP-seq data and *in planta* RNA-seq data indicates that chromatin modifications might contribute to the expression of pathogenicity related genes. *KMT1* and *KMT1/KMT6* deletions resulted in impaired *in vitro* growth and host colonization is consistently reduced compared to the WT while *KMT6* deletion only slightly reduced infection abilities. In all deletion backgrounds no large effect on gene expression was observed (Moeller *et al*., 2018). Analyses of gene expression in *KMT1* or *KMT6* mutants appear not to be the most appropriate approach to disentangle influence of histone modifications on the regulation of pathogenicity related genes. Therefore, we further investigated histone modification dynamics, *in planta*, in the WT strain. We investigated whether the up-regulation of putative effector genes at 13 dpi is associated with a difference in the histone methylation patterns between *in vitro* and *in planta* growth. We compared the histone methylation patterns in the genomic sequence of different genes that are lowly expressed *in vitro* and up-regulated during the lifestyle switch *in planta*, using ChIP-qPCR. The histone H4 encoding gene, constitutively expressed, and a transposable element, constitutively silenced (the DNA transposon, DTH_element 5_ZTIPO323, located on chromosome 9;) were used as controls in the experiment.

The three selected genes (Zt09_chr_5_00271; Zt09_chr_6_00192; Zt09_chr_9_00038) encode putative effectors among the 10 most expressed genes in *Z. tritici* at 13 dpi and these are considerably less expressed *in vitro* compared to the histone H4 gene (Table S8). The putative effector gene Zt09_chr_9_00038 encodes a hydrophobin-like protein considered to be a “core” necrotrophic effector, i.e. conserved in different isolates of *Z. tritici* (Haueisen *et al*., 2018). Using ChIP-qPCR, we confirm the *in vitro* association of the five genes with H3K4me2, H3K9me3 or H3K27me3 (“% of input” method). Consistently, the histone H4 encoding gene was associated with H3K4me2 while the TE sequence was associated with H3K9me3 and H3K27me3 and the three putative effector genes were located in heterochromatic domains (H3K9me3 or H3K27me3) *in vitro* (Figure 5; Table S8).

**Figure 5.**
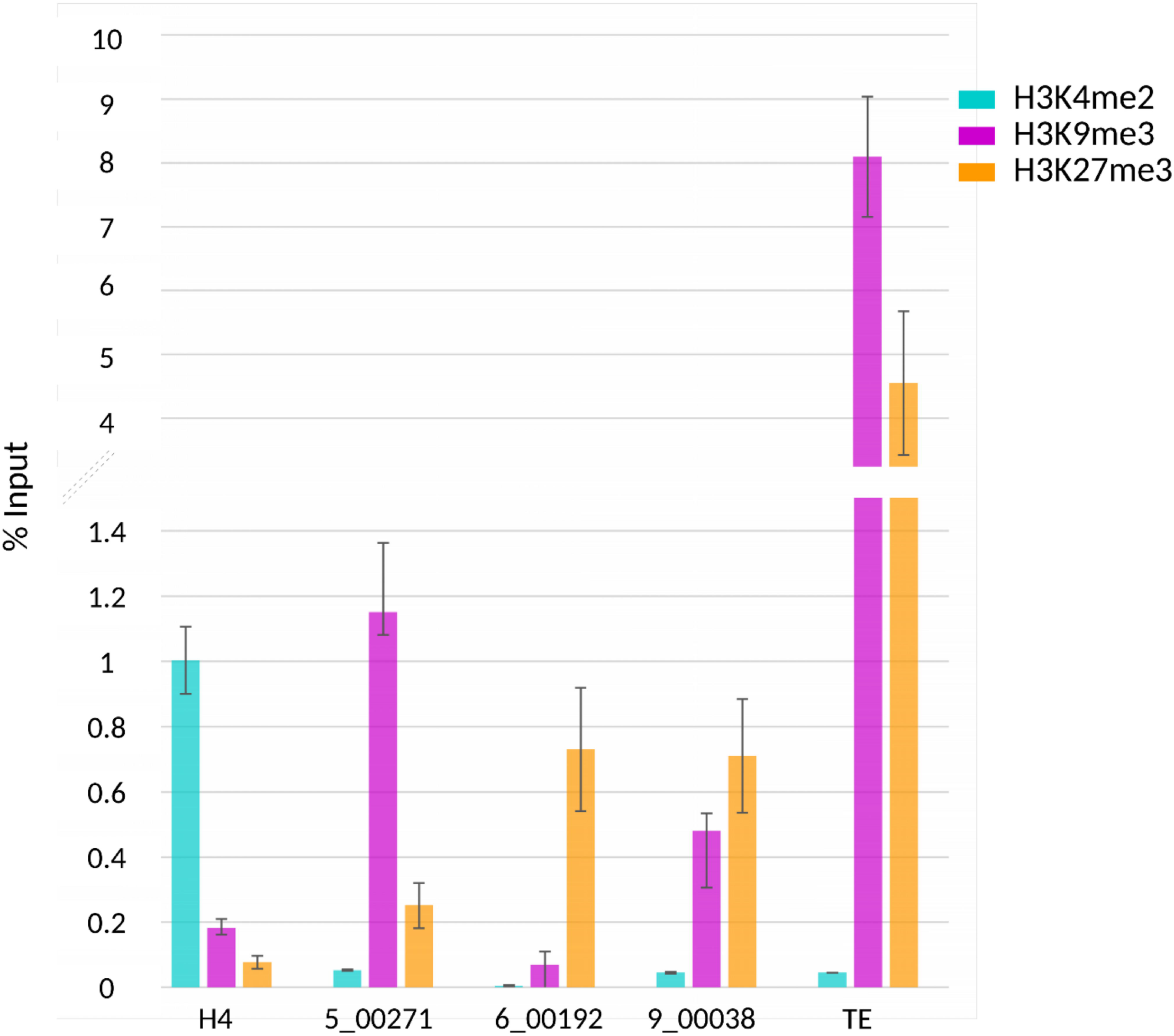
Putative effector genes are enriched in H3K9me3 and/or H3K27me3 *in vitro* while histone H4 is enriched in H3K4me2. Chromatin immunoprecipitation was performed during axenic culture to assess the levels of H3K4me2, H3K9me3 and H3K27me3 in the genomic sequence of four genes (H4, Histone H4: Zt09_chr_6_00256; 5_00271: Zt09_chr_5_00271; 6_00192: Zt09_chr_6_00192; 9_00038: Zt09_chr_9_00038) and a transposable element (DTH_element5_ZTIPO323).

For the *in planta* material, we confirmed the chromatin immunoprecipitation by a qPCR targeting the wheat Actin gene (Genebank accession number KC775780.1, cultivar BR34) at 13 dpi. We validated that there was no product for the wheat actin primers in the ChIP samples generated from fungal axenic culture. The abundance of H3K4me2 for the wheat actin gene was considerably higher compared to levels of H3K9me3 and H3K27me3 (Figure S1). In order to compare values obtained for the fungal genes *in vitro* and *in planta*, we calculated enrichment values of the targeted genes relatively to the fungal histone H4 gene. We firstly confirmed that this method led to the same conclusion as the “% input” method (Figure 5 and Table S8). Furthermore, we checked the difference in terms of enrichment between the gene promoter and the coding sequence: we applied qPCR following ChIP targeting the upstream gene sequence and the coding region of the gene Zt09_chr_6_00192. The levels of H3K4me2, H3K9me3, H3K27me3 shows the same profile across the promoter and coding sequence, i.e. level of any histone modifications is similarly high or low for the upstream gene sequence or the coding region of the gene (Figure 6) between the promoter and the coding sequence of a gene.

**Figure 6.**
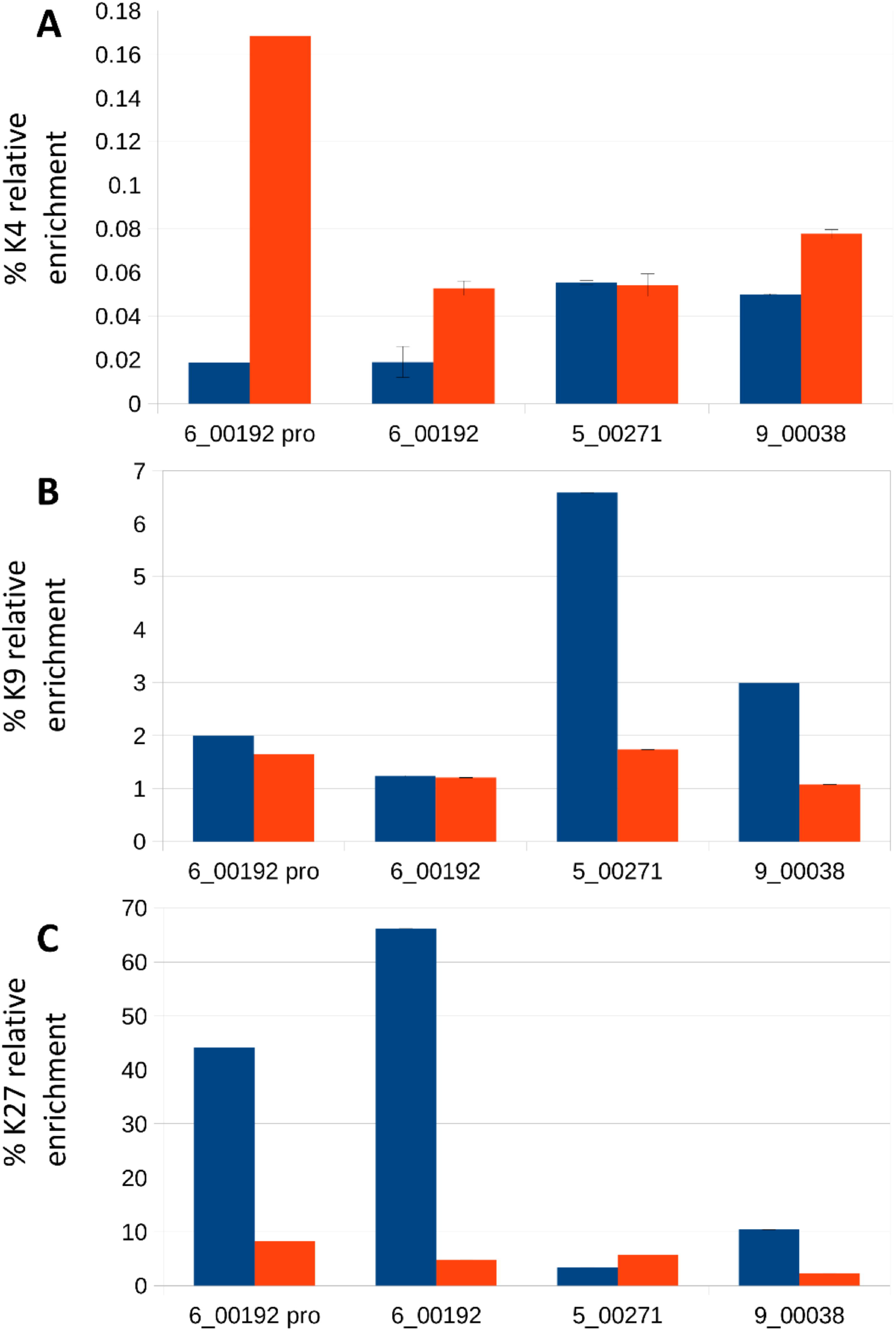
The up-regulation of three effector genes *in planta* is associated with a change of their histone modifications patterns. ChIP was performed at 13 days post infection of wheat leaves by *Zymoseptoria tritici* and relative enrichment of (A) H3K4me2, (B) H3K9me3 and (C) H3K27me3 was assessed using qPCR and compared during the axenic culture and at 13 dpi. 5_00271: Zt09_chr_5_00271; 6_00192: Zt09_chr_6_00192; 6_00192 pro: Zt09_chr_6_00192, promoter; 9_00038: Zt09_chr_9_00038; K4, K9 and K27: H3K4me2, H3K9me3 and H3K27me3 respectively. Blue: axenic culture; red: 13 dpi.

At 13 dpi, the three targeted effector genes are highly expressed (Table S8) and the relative enrichment of H3K4me2 remained low compared to histone H4 gene (% of H3K4me2 enrichment for these genes compared to histone H4 < 0.2). On the contrary, the levels of H3K9me3 decreased (a relative 3-fold lower amount of H3K9me3) for the genes Zt09_chr_5_00271 and Zt09_chr_9_00038. For two genes located in H3K27me3 *in vitro* (Zt09_chr_9_00038 and Zt09_chr_6_00192), levels of H3K27me3 were 6-22 fold lower at 13 dpi compared to axenic culture. This suggests that the up-regulation of these genes at 13 dpi is associated with a remodeling of the underlying chromatin structure in their genomic sequence (Figure 6) and that the influence of the chromatin structure on effector gene expression is due to a decrease of the heterochromatin marks rather than an increase of the euchromatic mark investigated here.

We have assessed the levels of H3K4me2, H3K9me3 and H3K27me3 for the TE relatively to the histone H4 gene: levels of H3K9me3 and H3K27me3 decreased compared to the levels *in vitro* but remained very high when compared to histone H4 and the three tested putative effector genes, which are up-regulated at this stage. Assessment of the chromatin state at the genomic loci of these genes confirms that constitutively expressed genes, such as the histone H4 gene, remain associated with the euchromatic modification H3K4me2 while the expression changes of the three effector genes tested are associated with changes of the associated histone modifications during wheat infection. This data provide evidence of an involvement of chromatin dynamics for the *in planta* regulation of secreted protein encoding genes putatively involved in pathogenicity.

## Discussion

In some plant pathogenic fungi, including *Z. tritici*, the expression profiles of effector genes was shown to be highly dynamic during host infection. By correlating genomic coordinates of predicted effector genes to transcriptome and epigenome data, we here provide evidence for a role of a chromatin-based gene regulation during plant infection of *Z. tritici*. In particular, candidate effector genes up-regulated during the transition from asymptomatic to necrotrophic growth are significantly enriched in regions of the genome that are transcriptionally silenced by a heterochromatin structure of the DNA during axenic growth. In this study, we analyzed levels of different histone modifications typical for heterochromatin or euchromatin *in planta* by applying ChIP-qPCR targeting the genomic loci of three effector genes that are among the most up-regulated genes at 13 dpi. Our experimental work provides evidence that histone modifications H3K9me3 and H3K27me3 play a role for the transcriptional regulation of putative effector genes in *Z. tritici* during wheat infection. Our targeted analysis *in planta* thereby suggests that the dynamic expression of effector genes can be associated with a dynamic of the heterochromatin marks H3K9me3 and H3K27me3.

### *Genes located in a heterochromatin domain* in vitro *are enriched in putative virulence related genes*

In other species, such as *L. maculans, Fusarium oxysporum*, Aspergilli species, genes located in regions that are TE-rich, subtelomeric, lineage-specific, are often enriched in genes encoding putative effectors or genes involved in secondary metabolite production (Rouxel *et al*., 2011; Grandaubert *et al*., 2014; Ma *et al*., 2010; Faino *et al*., 2016; Vlaardingerbroek *et al*., 2016). We find here that TE-rich genomic compartments of *Z. tritici* encode species-specific genes and genes encoding putative effector proteins. The fact that TE-rich regions are enriched in species-specific genes was also shown at the within species level (Plissonneau *et al*., 2016). Interestingly, while the *Z. tritici* reference strains presents eight accessory chromosomes which contain twice as much TEs as the core chromosomes, they are not enriched in putative secreted protein encoding genes as only nine are located on the accessory chromosomes. Genes encoding other types of pathogenicity related genes such as PKS and NRPS were also enriched in H3K27me3 domains of core chromosomes. Together, these data show that in *Z. tritici*, pathogenicity-related genes and genes involved in host specificity are enriched in repeat-rich, heterochromatic regions on the core chromosomes.

### *Genes highly expressed at the lifestyle switch towards necrotrophy are enriched in genes located in heterochromatin* in vitro *and in effector candidates*

Using our previous RNA-seq data generated *in vitro* and at 4 and 13 dpi (Kellner *et al*., 2014; Haueisen *et al*., 2018), we investigated patterns of gene expression as a function of their location in euchromatin or a heterochromatin domain *in vitro*. Genes over-expressed at 4 dpi compared to axenic culture and at 13 dpi compared to 4 dpi are enriched in putative pathogenicity-related genes. Interestingly, we found that genes highly expressed at the onset of the infection are not enriched in *in vitro* H3K4me2, H3K9me3 or H3K27me3 domains. However, genes highly expressed at the switch to necrotrophic growth are, *in vitro*, associated with TE sequences and H3K9me3 and H3K27me3. The different relevance of H3K9me3 and H3K27me3 at the two studied infection stages suggests that the pattern of histone modifications is dynamic during infection, and possibly influenced by signals from the host tissue and / or by fungal development *in planta*. It is possible that transcription of the genes highly expressed during the second wave of effector gene expression remains suppressed during early host infection to avoid induction of host immune responses. During early host infection, specific effectors are likely relevant to suppress the plant immune system upon host penetration and biotrophic fungal establishment.

Recently, functional analyses in *Z. tritici* have identified two genes encoding avirulence proteins (i.e., effector that can be recognized by the immune system of the plant activating defense reactions): Avr3D1 and AvrStb6, over-expressed during wheat infection just before the switch to necrotrophic growth (Zhong *et al*., 2017; Meile *et al*., 2018). In our ChIP-seq data, *AvrStb6*, located in a subtelomeric region, is associated with H3K9me3 and H3K27me3 while *Avr3D1*, located close to repetitive sequences, is associated with H3K27me3. Some of the genes encoding candidate necrosis inducing proteins, highly expressed at the switch to necrotrophy, identified in a previous study (Ben M’Barek *et al*., 2015) are also associated with heterochromatin domains *in vitro*, for instance genes Zt09_chr_5_00190 or Zt09_chr_2_01243. The association of these genes with H3K27me3 or TEs and their expression patterns support our findings drawn at the genome wide scale.

### *The histone modification patterns is dynamic between* in vitro *and* in planta *stages for a few effector genes, influencing their expression*

The role of the chromatin structure for the regulation of effector genes in TE-rich regions has been investigated in *L. maculans*, whereby RNAi-silenced transformants were generated for genes encoding HP1 and KMT1, two players involved in heterochromatin assembly and maintenance (Soyer *et al*., 2014). In this fungus, silencing of *HP1* and *KMT1* led to an up-regulation of genes located in TE-rich regions, notably effector genes, correlated with a decrease in H3K9me3 (Soyer *et al*., 2014). The same experimental strategy was applied to investigate the role of histone modifications for the regulation of secondary metabolite-encoding genes in other fungal species including *F. graminearum, E. festucae* and Aspergilli species (Reyes-Dominguez *et al*., 2012; Chujo and Scott, 2014; Gacek-Matthews *et al*., 2015; Gacek-Matthews *et al*., 2016). In *E. festucae*, ChIP-qPCR was also applied *in planta* to analyze histone modification patterns at the genomic loci of secondary metabolite gene clusters located in subtelomeric areas. In *Fusarium fujikuroi*, deletion of the H3K36 methyltransferase or *KMT6* resulted in deregulation (either up- or down-regulation) of some secondary metabolite gene clusters (Janevska *et al*., 2018; Studt *et al*., 2016). These studies, together with other analyzing gene expression at the genome-wide scale in plant pathogenic fungi (Connolly *et al*., 2013; Soyer *et al*., 2014; Moeller *et al*., 2018) show that loss of histone methyltransferases result in derepression of some genes associated with the histone modifications while a fraction remains silenced suggesting that a very complex regulatory network is involved in their transcriptional control. As in Soyer *et al*. (2014), our analysis of gene expression in *KMT1, KMT6* mutants of *Z. tritici* (Moeller *et al*., 2018; our study) has shown that pathogenicity-related encoding genes and genes expressed upon infection, are enriched within the deregulated genes due to at least one of the deletion. In order to analyze influence of histone modifications in the WT context and during plant infection, we applied for the first time ChIP-qPCR *in planta* to analyze levels of H3K4me2, H3K9me3 and H3K27me3 of three genes encoding effector candidates which are highly expressed at 13 dpi, as well as the constitutively expressed histone H4 gene and a transposable element sequence. We could demonstrate that levels of H3K4me2, H3K9me3 and H3K27me3 for each targeted loci correlated with their expression *in vitro* and at 13 dpi. For the repetitive element, levels of H3K9me3 and H3K27me3 were high *in vitro* as well as *in planta* likely reflecting the efficient silencing of TEs in some regions of the genome. However, effector genes associated with a high level of H3K9me3 and H3K27me3 *in vitro* showed significantly reduced levels of these marks *in planta* consistent with their strong up-regulation. This indicates that the chromatin structure is dynamic between axenic culture and different infectious stages and instrumental for regulation of gene expression, notably of at least some effector genes in *Z. tritici*.

In conclusion, our study indicates a prominent role of chromatin-based gene regulation during wheat infection of *Z. tritici*. The close association of putative effector genes with repeat-rich DNA may facilitate rapid evolution of these genes. Furthermore, the association with heterochromatin in these genomic compartments provides variability at the transcriptional level possibly further facilitating the defeat of host defenses by this important wheat pathogen.

## Experimental Procedures

### Fungal strain and plant cultivar

The *Z. tritici* isolate Zt09 was used throughout the study. Zt09 is derived from the reference isolate IPO323, and differs by the absence of chromosome 18, lost during *in vitro* culture (Kellner *et al*., 2014). Cultures were grown on solid yeast malt sucrose (YMS) agar (4 g yeast extract, 4 g malt extract, 4 g sucrose, 20 g bacto agar, 1 liter H_2_O) at 18°C in the dark. For the *in planta* experiments conducted in this study, *Triticum aestivum* cultivar Obelisk (Wiersum Plantbreeding, Winschoten, the Netherlands) was used. The *kmt1, kmt6* and *kmt1/kmt6* mutants were generated by Moeller *et al*. (2018).

### Identification of genes associated with specific histone modifications

The annotation of the *Z. tritici* isolate presented in Grandaubert *et al*. (2015) was used in this study including genes predicted to encode putative effectors and Polyketide Synthases/NonRibosomal Peptide Synthetases. To distinguish genes associated with the different histone marks H3K4me2, H3K9me3 and H3K27me3, we used previously generated *in vitro* ChIP-seq datasets (Schotanus *et al*., 2015) available under the SRA accession number SRP059394. The Integrative Genome Viewer (IGV; http://www.broadinstitute.org/software/igv; Thorvaldsdóttir *et al*., 2013) was used to visualize location of each domains along the genome. Statistically enriched regions were identified with RSEG (Song and Smith, 2011). We determined the *in vitro* chromatin state of each annotated gene using the previously published chromatin maps (Schotanus et *al*., 2015). Genes were considered to be associated with a given post-translational histone modification when partially (at least one bp) or completely located in a H3K4me2, H3K9me3 or H3K27me3 domain. A Chi^2^ test was applied to identify statistical enrichment of euchromatic or heterochromatic domains for certain categories of genes: the expected proportion of a given category of genes across the entire genome was compared to the observed distribution of the gene category in the H3K4me2, H3K9me3 or H3K27me3 domains. Enrichment was considered significant with a *P* value < 0.01. All analyses were done in R (www.r-project.org).

### RNA-sequencing datasets

To correlate gene expression with location in a given chromatin domain during *Z. tritici* infection of wheat, we used previously generated *in vitro* and *in planta* (four and 13 dpi) RNA-seq datasets of Zt09 (Kellner *et al*., 2014; Haueisen *et al*., 2018). RNA-seq data from the *in vitro* grown cultures and the *in planta* stages are accessible at the NCBI Gene Expression Omnibus respectively through accession number GSE54874 and GSE106136 (Kellner *et al*., 2014; Haueisen *et al*., 2018). RNA-seq data from the *kmt1, kmt6* and *kmt1/kmt6* mutants during *in vitro* growth were generated by Moeller *et al*. (2018) and at Sequence Read Archive under BioProject ID PRJNA494102.

RPKM values for each gene in a given condition were estimated using Cufflinks (Trapnell *et al*., 2010) as already described in Kellner *et al*. (2014). The total read counts per transcript were estimated in HTSeq using the intersection-strict mode (Anders *et al*., 2015) and genes differentially expressed were identified with the EdgeR package (Robinson *et al*., 2010) with a log2 Fold Change ≤ −1 or ≥ 1 and an associated false discovery rate ≤ 0.001 (McCarthy *et al*., 2012). As previously conducted in a study of gene expression in *Z. tritici* (Kellner *et al*., 2014), we only included genes with an RPKM value ≥ 2 in the condition in which the gene is up-regulated.

### Plant experiments for chromatin immunoprecipitation quantitative PCR analyses

Plant infections were done as previously described (Habig *et al*., 2017). In brief, the second leaves of wheat seedlings were inoculated with a solution containing 1.10^7^ cells/ml supplemented with 0.1% Tween by brushing an area of seven cm after 11 days of seedling growth. Plant material was harvested four and 13 dpi for ChIP-qPCR. As a control, non-inoculated leaves were also harvested four and 13 dpi.

### Chromatin immunoprecipitation (ChIP) and quantitative PCR (qPCR)

ChIP with antibodies against H3K4me2, H3K9me3 and H3K27me3 was performed on cells from axenic cultures as previously described (Soyer *et al*., 2015). For *in planta* ChIP, 12 infected leaves were harvested (i.e. three leaves and four technical replicates) at 13 dpi and the same protocol was used except that “native ChIP” (without formaldehyde crosslinking) was applied from 100 mg of infected plant material. All DNA extractions were done with the Wizard^®^ SV Gel and PCR Clean Up system kit (Promega, Madison, USA).

Based on our analyses of transcriptome, genome and ChIP-seq data, we selected three candidate genes for ChIP-qPCR analyses: Zt09_chr_5_00271, Zt09_chr_6_00192, Zt09_chr_9_00038. These genes are lowly expressed *in vitro* and among the 10 most expressed genes in *Z. tritici* 13 dpi of wheat. Moreover, they are associated with heterochromatin during *in vitro* growth (Table S8; Schotanus *et al*., 2015). Furthermore, we included the sequence of a transposable element (DTH_element5_ZTIPO323, located on chromosome 9, position 29,065-30,735) as well as the gene encoding histone H4 (ID Zt09_chr_6_00256) as controls for precipitation with anti-H3K9me3, anti-H3K27me3 antibodies and anti-H3K4me2 antibody, respectively.

Quantitative PCR (qPCR) was performed with SYBR Green PCR Master Mix (Applied Biosystem, Foster City, USA) on a 7900 Real Time PCR System (Applied Biosystems). Two biological replicates, and two technical replicates were processed. Primers were designed with the OligoPerfect Designer (ThermoFisher Scientific) to target amplification of products between 80 and 150 bp (Table S9). The efficiency of PCR primers was verified on genomic DNA as described previously (Taylor *et al*., 2010). For RT-qPCR, Ct values were analyzed as described in Muller *et al*. (2002) for analysis of expression profiles and the Glyceraldehyde-3-phosphate dehydrogenase (GAPDH)-encoding gene (ID Zt09_chr_2_00354) was used as a constitutively expressed reference gene, as in Poppe *et al*. (2015). A positive control for the *in planta* ChIP experiment was obtained by qPCR amplification of the wheat actin gene (Table S9). For qPCR after *in vitro* ChIP, the immunoprecipitated fraction of each gene was calculated by the “% of input” method (Lin *et al*., 2012). Following the qPCR experiment, the Ct values (number of cycles required for the fluorescent signal to cross the threshold) were retrieved from the 7900 SDS software. To compare enrichment of the histone modifications H3K4me2, H3K9me3, and H3K27me3 for each target gene *in vitro* and *in planta*, we compared Ct values of the target genes to Ct values of a reference gene, histone H4 (i.e., for example, ΔCtH3K4me2 = Ct gene H3K4me2 − Ct histone H4 H3K4me2). The same calculation was applied for the histone H4 gene, therefore ΔCt = 1 for each modification for this gene. Finally, the enrichment of each target gene with the three histone modifications was calculated as described by Lin *et al*. (2012) using %H3K4me2 = 100/2^ΔCt^. This enrichment was compared for each gene *in vitro* and *in planta* (at 13 dpi).

## Acknowledgments

The authors thank members from the “Environmental Genomics” group, in particular Mareike Moeller, and J.Y. Dutheil, for fruitful discussion. J.L. Soyer was funded by a Young Scientist Funding by INRA. The study was supported by a Max Planck fellowship to EHS.

## Author contributions

Conceived and designed the experiments: JLS, JG, EHS. Acquisition, analysis or interpretation of the data: JLS, JG, JH, KS, EHS. Writing of the manuscript: JLS, EHS.

## Supplementary data legends

**Figure S1. Wheat actin was enriched in histone H3K4me2 and was used as a control for chromatin immunoprecipitation *in planta***. ChIP was performed at 13 days post infection of wheat leaves by *Zymoseptoria tritici*. Enrichment of the wheat actin gene in H3K4me2, H3K9me3 and H3K27me3 was assessed using qPCR.

**Table S1. Enrichment analysis of PKS/NRPS- and orphan genes in a H3K4me2, H3K9me3, H3K27me3 or H3K9me3/H3K27me3 domain *in vitro***.

^a^Genes encoding PKS, NRPS, and orphan genes as predicted by Grandaubert *et al*. (2015);

^b^Genes with a RPKM ≥ 2;

^c^Genes with Log2 Fold Change (RPKM) ≥ 1 and FDR ≤ 0.001 at 4 days post infection (dpi) compared to axenic culture or 13 dpi compared to 4 dpi were considered as up-regulated at a given time point compared to the other. Only the genes with a RPKM ≥ 2 at least at the time point during which it is up-regulated were kept for the analysis;

^d^Genes with Log2 Fold Change (RPKM) ≤ −1 and FDR ≤ 0.001 at 4 days post infection (dpi) compared to axenic culture or 13 dpi compared to 4 dpi were considered as down-regulated at a given time point compared to the other;

^e^Genes located in a H3K4me2, H3K9me3, H3K27me3 or H3K9me3/H3K27me3 *in vitro* as identified with RSEG (see Experimental Procedures; Schotanus *et al*., 2015).

Blue: the given domain contains significantly less genes of the given category compared to the rest of the genome; orange: the given domain is enriched for the category of genes compared to the rest of the genome.

**Table S2. Enrichment analysis of genes of a given category in a H3K4me2, H3K9me3, H3K27me3 or H3K9me3/H3K27me3 domain *in vitro***.

^a^Genes of “unknown function” are genes encoding predicted or hypothetical proteins; genes encoding CAZymes as predicted by Grandaubert *et al*. (2015);

^b^Genes located up to 2 kb upstream or downstream to a transposable element sequence as predicted by Grandaubert *et al*. (2015);

^c^Genes with a RPKM ≥ 2;

^d^Genes with Log2 Fold Change (RPKM) ≥ 1 and FDR ≤ 0.001 at 4 days post infection (dpi) compared to axenic culture or 13 dpi compared to 4 dpi were considered as up-regulated at a given time point compared to the other. Only the genes with a RPKM ≥ 2 at least at the time point during which it is up-regulated were kept for the analysis;

^e^Genes with Log2 Fold Change (RPKM) ≤ −1 and FDR ≤ 0.001 at 4 dpi compared to axenic culture or 13 dpi compared to 4 dpi were considered as down-regulated at a given time point compared to the other;

^f^Genes located in a H3K4me2, H3K9me3, H3K27me3 or H3K9me3/H3K27me3 *in vitro* as identified with RSEG (see Experimental Procedures; Schotanus *et al*., 2015).

Blue: the given domain contains significantly less genes of the given category compared to the rest of the genome; orange: the given domain is enriched for the category of genes compared to the rest of the genome.

**Table S3. PFAM analysis of the genes located in H3K4me2 domains *in vitro***.

**Table S4. PFAM analysis of the genes located in H3K9me3/H3K27me3 domains *in vitro***.

**Table S5. PFAM analysis of the genes located in H3K9me3 domains *in vitro***.

**Table S6. PFAM analysis of the genes located in H3K27me3 domains *in vitro***.

**Table S7. Enrichment analysis of putative effector genes in the vicinity of a transposable element or in a H3K4me2, H3K9me3, H3K27me3 and H3K9me3/H3K27me3 domain *in vitro*, and according to their expression**.

^a^Genes encoding predicted secreted proteins (SP) were considered as genes encoding putative effector genes for this study, as predicted by Grandaubert *et al*. (2015);

^b^Genes with Log2 Fold Change (RPKM) ≥ 1 and FDR ≤ 0.001 at 4 days post infection (dpi) compared to axenic culture or 13 dpi compared to 4 dpi were considered as up-regulated at a given time point compared to the other. Only the genes with a RPKM ≥ 2 at least at the time point during which it is up-regulated were kept for the analysis;

^c^Genes with Log2 Fold Change (RPKM) ≤ −1 and FDR ≤ 0.001 at 4 dpi compared to axenic culture or 13 dpi compared to 4 dpi were considered as down-regulated at a given time point compared to the other;

^d^Genes located up to 2 kb upstream or downstream to a transposable element sequence as predicted by Grandaubert *et al*. (2015);

^e^Genes located in a H3K4me2, H3K9me3, H3K27me3 or H3K9me3/H3K27me3 *in vitro* as identified with RSEG (see Experimental Procedures; Schotanus *et al*., 2015).

Blue: the given domain contains significantly less genes of the given category compared to the rest of the genome; orange: the given domain is enriched for the category of genes compared to the rest of the genome.

**Table S8. Trimethylation of lysine 9 and / or 27 of histone H3 is reduced for three candidate effector genes up-regulated 13 days post infection of wheat leaves compared to axenic culture**.

^a^Orphan genes as described by Grandaubert *et al*. (2015);

^b^Genes located at a distance below 2 Kb of a transposable element (Grandaubert *et al*., 2015);

^c^Relative enrichment, *in vitro* or 13 dpi, of H3K4me2, H3K9me3 or H3K27me3 for the given locus compared to histone H4 encoding gene (Zt09_chr_6_00256);

^d^Gene expression *in vitro* (Kellner *et al*., 2014);

^e^Gene expression from Haueisen *et al*. (2018);

^f^Expression rank of the candidate genes at 13 days post infection (dpi).

**Table S9. List of primers used in this study**.

