## Supplementary figures and images for "*In planta* chromatin immunoprecipitation in *Zymoseptoria tritici* reveals chromatin-based regulation of putative effector gene expression"

### Supplemental Figure 1

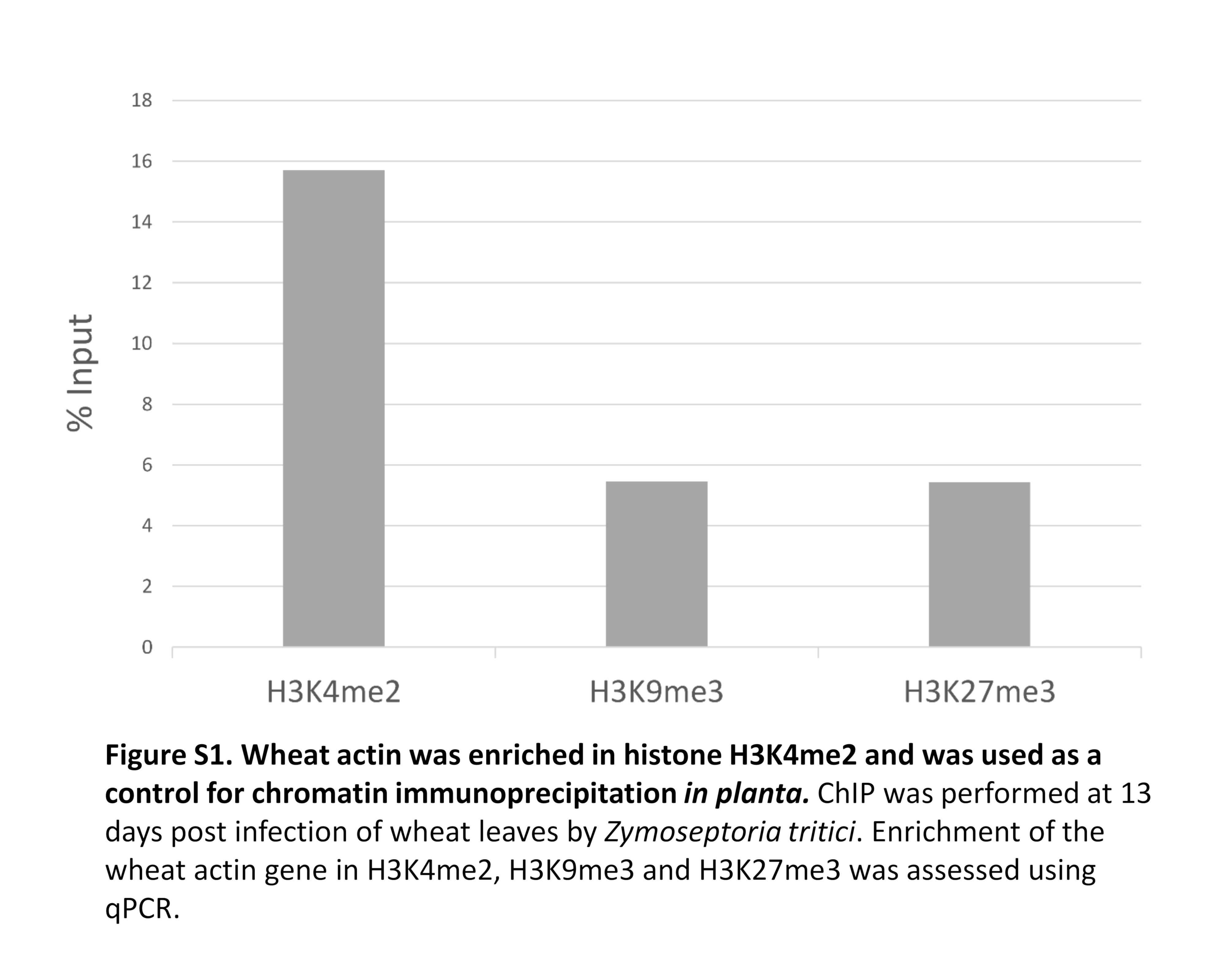
